# A Whole-Brain 3D Myeloarchitectonic Atlas: Mapping the Vogt-Vogt Legacy to the Cortical Surface

**DOI:** 10.1101/2022.01.17.476369

**Authors:** Niels A. Foit, Seles Yung, Hyo Min Lee, Andrea Bernasconi, Neda Bernasconi, Seok-Jun Hong

**Affiliations:** Neuroimaging of Epilepsy Laboratory, McConnell Brain Imaging Center, Montreal Neurological Institute, Montreal, QC, Canada; Department of Neurosurgery, Medical Center - University of Freiburg, Freiburg, Germany; Center for Neuroscience Imaging Research, Institute for Basic Science, Suwon, Korea; Department of Biomedical Engineering, Sungkyunkwan University, Suwon, Korea; Center for the Developing Brain, Child Mind Institute, NY, USA

**Keywords:** Myeloarchitecture, MNI space, intracortical depth profiling, FreeSurfer

## Abstract

Building precise and detailed parcellations of anatomically and functionally distinct brain areas has been a major focus in Neuroscience. Pioneer anatomists parcellated the cortical manifold based on extensive histological studies of post-mortem brain, harnessing local variations in cortical cyto- and myeloarchitecture to define areal boundaries. Compared to the cytoarchitectonic field, where multiple neuroimaging studies have recently translated this old legacy data into useful analytical resources, myeloarchitectonics, which parcellate the cortex based on the organization of myelinated fibers, has received less attention. Here, we present the neocortical surface-based myeloarchitectonic atlas based on the histology-derived maps of the Vogt-Vogt school and its 2D translation by Nieuwenhuys. In addition to a myeloarchitectonic parcellation, our package includes intracortical laminar profiles of myelin content based on Vogt-Vogt-Hopf original publications. Histology-derived myelin density mapped on our atlas demonstrate close overlap with *in vivo* quantitative MRI markers for myelin and relates to cytoarchitectural features. Complementing the existing battery of approaches for digital cartography, the whole-brain myeloarchitectonic atlas offers an opportunity to validate imaging surrogate markers of myelin in both health and disease.

**Highlights:**

- Our myeloarchitectonic atlas builds on extensive meta-analyses-derived and ground-truth histological data.
- Our atlas provides qualitative and quantitative 3D information on cortical myelin architecture.
- MRI surrogate markers of myelin demonstrate close overlap with histological cortical parcellations, supporting biological validity of non-invasive metrics.
- This atlas can be seamlessly integrated into widely used neuroimaging analysis software to inform studies in health and disease.

## Introduction

Obtaining precise and detailed parcellations of anatomically and functionally distinct brain areas has been the focus of Neuroscience research for over a century (Zilles et al., 2015; Zilles and Amunts, 2010). Among neuroanatomists of the early 20^th^ century, Brodmann, Vogt and Vogt, and von Economo and Koskinas ardently worked towards generating highly detailed histological maps of the human cortex based on post-mortem data (Triarhou, 2007; Zilles and Amunts, 2010). To parcellate the neocortex, they relied on cytoarchitectonics, characterizing size, shape and distribution of cell bodies across cortical layers (Amunts et al., 2005; Smith, 1927; Triarhou, 2007), and myeloarchitectonics, which studies layering, arrangement, packing and density of myelinated fibers and bundles (Batsch, 1955; Hopf, 1968; Nieuwenhuys, 2013; Strasburger, 1937; Vogt and Vogt, 1919). These histology-derived maps have set the basis for MRI-derived *in vivo* parcellations of cortical boundaries (Huntenburg et al., 2017; Mendes et al., 2019; Nieuwenhuys and Broere, 2020). Nevertheless, knowledge on cytoarchitectonics today remains heavily influenced by the classic work of Brodmann (Brodmann, 1907; Zilles and Amunts, 2010) and many contemporary neuroimaging toolkits contain modified versions of Brodmann’s seminal map (Eickhoff et al., 2005; Talairach et al., 1993).

Compared to the cytoarchitectonic field, myeloarchitectonics has received less attention. This may be due to the paucity of histology-based myeloarchitectonic data to establish the biological substrates of several MRI markers for myelin characteristics (Lazari and Lipp, 2021; van der Weijden et al., 2020), such as quantitative T1 mapping (Lutti et al., 2014; Marques et al., 2017) and neurite orientation and dispersion density imaging (Zhang et al., 2012). Recently, Nieuwenhuys and co-workers (Nieuwenhuys and Broere, 2017) mapped Vogts myeloarchitectonic atlases to a non-digital, 2D representation of the Montreal Neurological Institute (MNI) Colin27 brain template, by means of manual translations from paper-embedded figures. Harnessing the data generated by Hopf, they further integrated area-specific myelin fiber density estimates into their myeloarchitectonic map (Hopf, 1968, 1957, 1956, 1955; Nieuwenhuys and Broere, 2017).

While 2D maps render the knowledge from historical postmortem data more accessible, they cannot be used in quantitative neuroimaging analyses, which usually require 3D stereotaxic volumetric data representation or cortical surface formats. In an effort to address this gap, we built a 3D myeloarchitectonic atlas (MYATLAS) in common space, translating the Nieuwenhuys’ boundaries of the Vogt-Vogt atlas to the Colin27 brain template and standard cortical surfaces. Besides providing a *ready-to-use* myeloarchitectonic parcellation, we generated intracortical laminar profiles of myelin content from the photometric data gathered from Vogt-Hopf publications. To validate our atlas, we quantified the similarity between the histology-based myelin density and profiles from *in vivo* MRI markers obtained from MP2RAGE-derived qT1 mapping (21, 34, 35). Moreover, we integrated cytoarchitectonic features (Scholtens et al., 2018). Finally, to facilitate integration into existing image processing pipelines, the proposed myeloarchitectonic atlas together with source codes are made publicly available.

## Materials and Methods

After a short review of prior work, sections below detail the methodology used for the creation of our 3D myeloarchitectonic atlas (MYATLAS) and necessary processing steps to obtain cortical depth profiles from Vogt-Vogt myeloarchitectonic parcellations, as well as cross-modal correlations between myelo- and cytoarchitectural features and *in vivo* qT1 mapping. We further provide instructions for registering the MYATLAS to individual MRI data.

### Summary of prior work

Nieuwenhuys and co-workers recently performed a meta-analysis on potential usability of the historical myeloarchitectural data aggregated from Vogts’ papers (see Nieuwenhuys and Broere, 2017 for details). The original cortical division by the Vogts school consisted of a total of 180 myeloarchitectonic cortical fields (64 frontal, 30 parietal, 63 temporal, 17 occipital, and 6 insular areas). Their associates Hopf and Vitzthum further refined this work by subdividing the parietal and occipital lobes into multiple sub-areas (Hopf, 1957, 1956, 1955), resulting in 214 regions (64 frontal, 60 parietal, 63 temporal, 21 occipital, and 6 insular). Nieuwenhuys et al. applied semi-automatic topological translations and boundary averaging across 17 different views of this analog dataset to project myeloarchitectonic parcellations onto the MNI-Colin27 single subject brain in a non-digital figure format (**Figure 1**).

**Figure 1.**
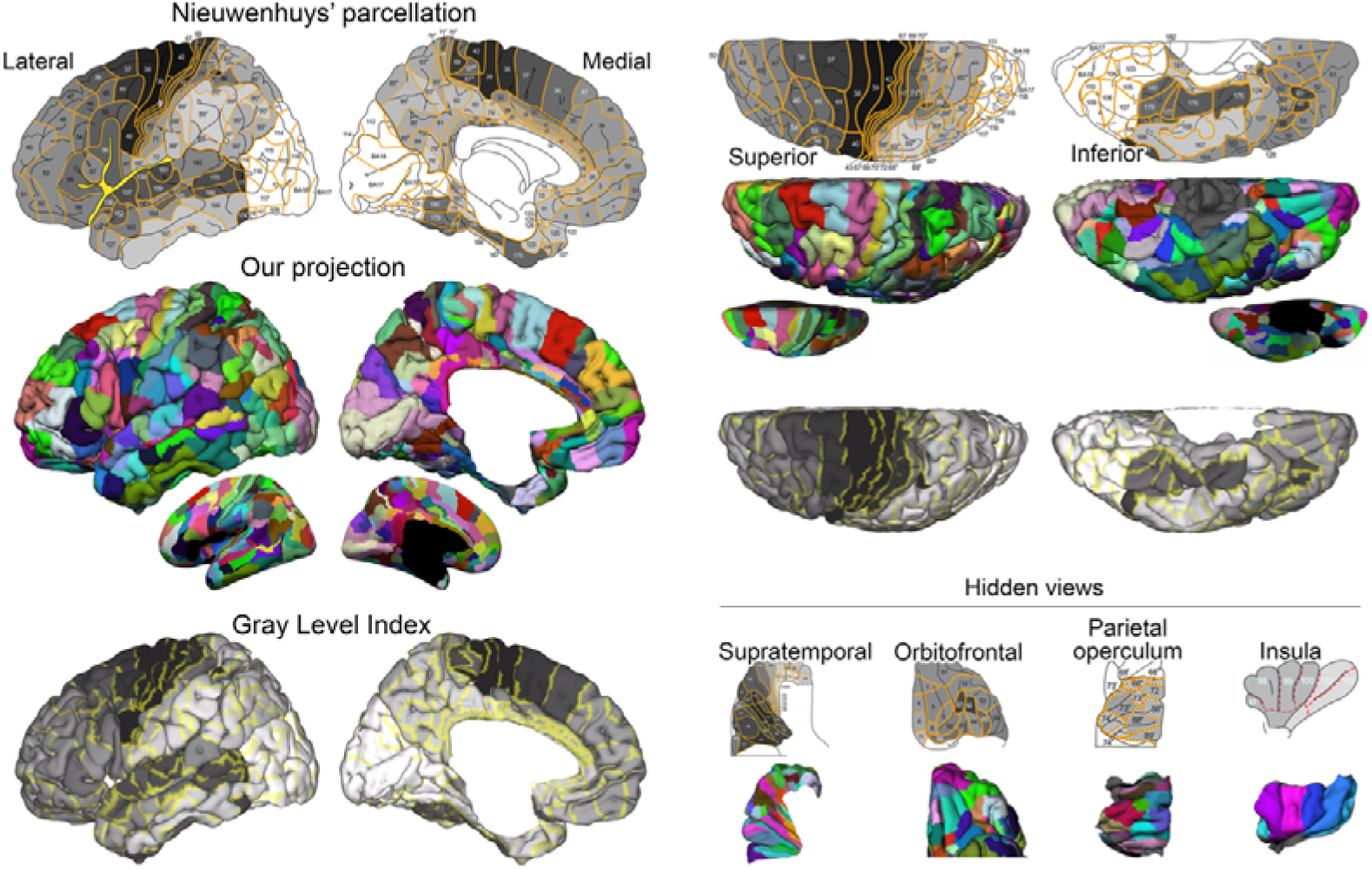
Construction of the 3D myeloarchitectonic MNI atlas (MYATLAS). Manual cortical parcellations (based on the topological transformation of Vogt-Vogt parcellation) across view planes (lateral, medial, superior, inferior; on pial and inflated surfaces), with the 3D projection and mean gray level index measuring the degrees of myelination. Hidden areas within the orbitofrontal region, the supratemporal lobe, and the parietal operculum were labeled by extracting view planes similar to the original publication; they were then merged back with the whole-brain surface once segmentations were finished (panel on the right bottom).

In addition to boundaries, the Vogts investigated intracortical penetration patterns of tangential and radial fiber bundles across cortical laminae (Vogt and Vogt, 1919). They defined major categories of myeloarchitectonic profiles based on two criteria (**Figure 2A**). The first criterion related to the presence of transverse, densely myelinated cortical layers (the bands of Baillarger). Accordingly, cortical specimens can be classified into 4 categories: *bistriate* (two horizontal myelin-rich bands), *unistriate* and *unitostriate* (both indicating only one visible band; for the former, a single cortical layer is covered, whereas for the latter multiple layers are covered by the band), and *astriate* (no striation). Notably, each category has a subtype, depending on the demarcation of a band boundary (*propebistriate*: barely recognizable bands of Baillarger*; propeunistriate*: ill-defined border of inner stripe*; propeastriate*: slight decrease of density in 5a/6a). The second criterion relates to the intrusion depth of the radiate bundles across cortical laminae, further classifying the cortex into *euradiate* (bundles reaching upper border of layer 3b), *infraradiate* (reaching upper border of layer 5b), and *supraradiate* (extending into layers 1-2).

**Figure 2.**
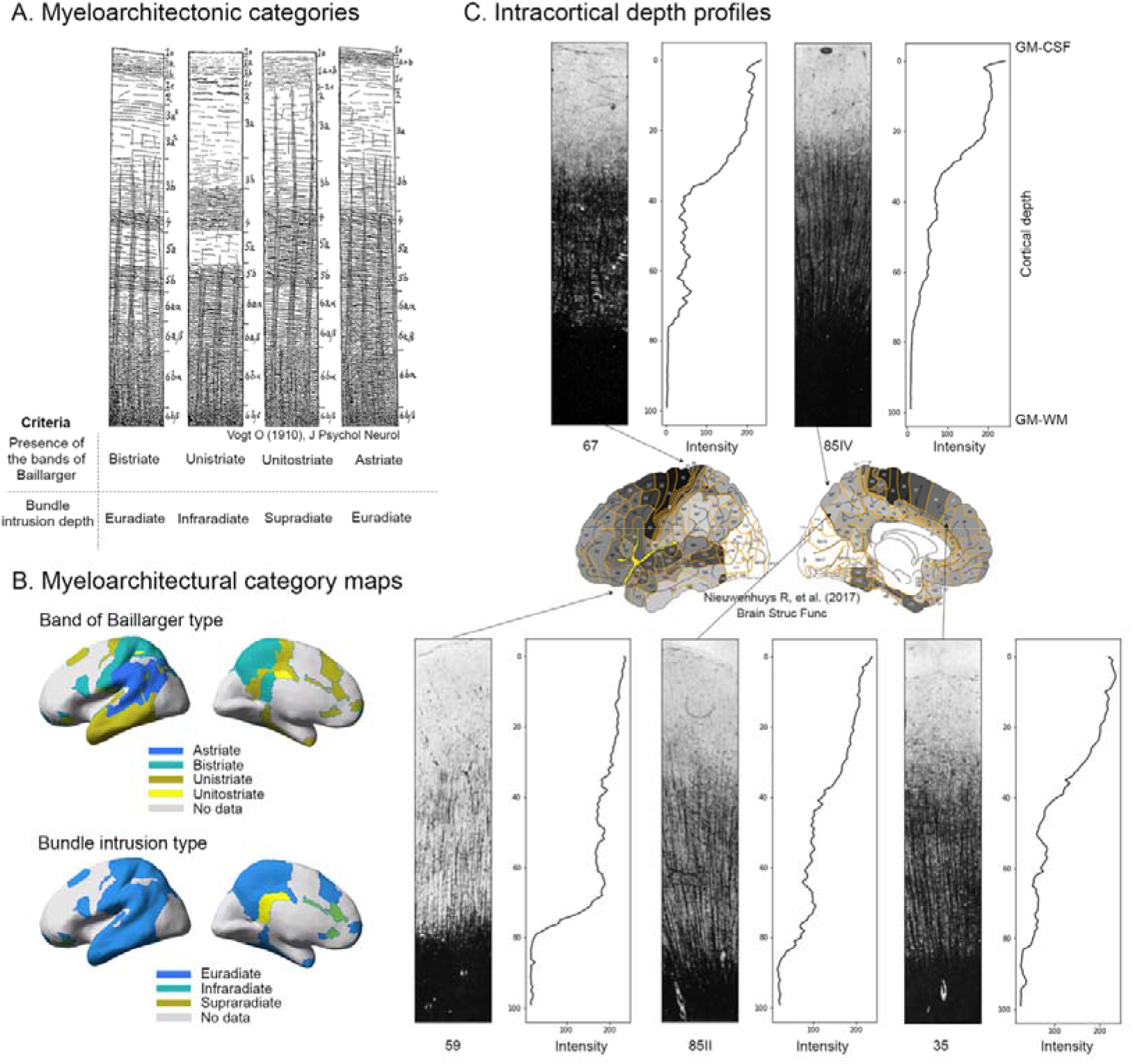
Intracortical depth profiling based on Vogt-Hopf histological data. **A)** Categories of myeloarchitectonic features based on laminar and bundle intrusion patterns; **B)** Mapping of feature information onto the cortical surface; *no data* indicate subtype information unavailable from the original literature. **C)** Examples of myelin-stained cortical areas across full cortical depth with their corresponding depth profiles (maps shown in **Supplementary Material 1**).

Following this work, Hopf (Hopf, 1968, 1957, 1956, 1955) further advanced these myeloarchitectonic parcellation by systematically recording frontal, parietal and temporal lobe myelin contents using analog photodensitometry. Notably, each region has distinct light absorption curves, or cortical depth profiles. This data was recently digitized by Nieuwenhuys (Nieuwenhuys and Broere, 2017), who introduced *mean grey levels* (MGL), *i.e*., digital quantifications of myelin fiber density per cortical area. Ranging from 0 to 255, lower values indicate densely myelinated areas, while high MGL are found in lightly myelinated areas (Edwards et al., 2018; Nieuwenhuys and Broere, 2017). Since MGL are only available for the frontal, parietal and temporal lobes, remaining areas were assigned 255 as a default value. Since Hopf based his work exclusively on the right hemisphere, MGL were available only for the right hemisphere in Nieuwenhuys’ work (Nieuwenhuys and Broere, 2017).

### MRI processing

We created a stereotaxic average of the individual Colin27 brain MRIs, comprising 27 T1 weighted scans with 1mm isotropic voxel resolution (Holmes et al., 1998). We then extracted 3D cortical surface models from this template using FreeSurfer (Fischl, 2012). Briefly, processing steps included gradient non-uniformity correction (Jovicich et al., 2006), registration to MNI stereotaxic space, intensity normalization, skull stripping, and segmentation into tissue classes (Fischl et al., 2004). Gray-white and gray-CSF interface models were generated through triangular surface tessellation yielding 163,842 vertices (Dale et al., 1999), followed by topology correction, inflation, and spherical registration to fsaverage (Fischl et al., 2001).

### 2D-to-3D translation of myeloarchitectural parcellations

As per previous procedures (Pijnenburg et al., 2021; Scholtens et al., 2015), the original Nieuwenhuys’ illustration (Nieuwenhuys et al., 2015) was split into eight view planes (lateral, medial, superior, inferior, orbitofrontal, supratemporal, parietal opercular, insular). Carefully cross-referencing different views, a single rater (SY) labeled each parcellation by comparing geometric landmarks between Nieuwenhuys ‘ illustrations and the convexity of the 3D Colin27 brain surface. Labelling was performed with “*tksurfer*” (surfer.nmr.mgh.harvard.edu/fswiki/TkSurfer). Notably, identifiable sulci on the original 2D Colin27 map such as the central and superior temporal sulcus served as systematic landmarks. To optimize anatomical matching of area boundaries, we further relied on the main sulcal patterns surrounding each region to be labeled. A second rater (SJH) evaluated accuracy of each label. For ambiguous areas, *i.e*., either a mismatch across view planes, or between 2D illustrations and 3D cortical surface, an inter-rater consensus on boundaries was reached to minimize discrepancies by carefully reviewing the original publications and applying manual corrections, if necessary (**Supplementary Table 1**).

An inherent limitation of Vogt-Vogt histological data is that all results were reported on the convex pial surface, which does not reveal buried sulci. We thus labelled areas located within these sulci on the white matter surface view on which they are clearly visible and intra-sulcal boundaries can be easily delineated at their bottom. To visualize areas hidden in the depths of the Sylvian fissure (*i.e*., insula, supratemporal lobe, parietal operculum), invisible even on the white matter surface, we extracted patches of their surfaces, delineated label boundaries and merged them back to the whole-brain data. Notably, some areas appeared multiple times across view planes, preventing their segmentation into a single coherent label. This was addressed by dividing areas into sub-regions; for example, for area 111 appearing differently in the lateral and medial views, we divided it into 111-l, 111-m, ‘1’ and ‘m’ refer to medial and lateral, respectively (see **Supplementary Table 1**). Finally, resulting labels were numbered in accordance to Vogt’s numeric convention and merged with color tables to create a single Freesurfer annotation file (.annot), which contains a total of 214 parcellations (**Figure 1C**).

### Quantitative cortical myelin content and intracortical depth profiling

MRI allows for *in vivo* quantification of myelin content of the cortical manifold (Stüber et al., 2014; Waehnert et al., 2014). However, validation requires access to histology, which has not been available in a digital format. We thus generated histology-derived quantitative myelin data by recording the MGL index (the averaged myelin fiber density) for each cortical field (Nieuwenhuys and Broere, 2017) and created a *ready-to-analyze* look-up table in excel format for the use with the MYATLAS (**Supplementary Material 1**, “Myeloarchitectural_table.xslx”). As the MGL for the insula and occipital lobes (34 parcels) were unavailable, they were omitted, totaling 187 values (64 frontal, 60 parietal and 63 temporal areas; **Supplementary Material 1**). We then mapped the myelination density (MGL values) onto the MYATLAS (**Figure 1**).

To extract myelin laminar depth profiles from histologically-stained microphotographs (Batsch, 1955; Brockhaus, 1940; Hopf, 1957, 1956, 1955; Strasburger, 1937; Vogt and Vogt, 1919), we screen-captured the histology photos with a fixed format and size, and estimated the gray level intensity across cortical laminae as a surrogate of myelin density. The digitized histological figures were normalized to make intensity values across photos comparable. Absolute gray values were then extracted and plotted as a normalized depth profile across all cortical layers (**Figure 2B**). Information on myeloarchitectonic features, such as fiber bundle types and layer-specific density of each cortical stain, were also recorded (**Supplementary Material 1, Figure 2C**).

### Correlation between myeloarchitectonic features and in-vivo myelin proxy data

We cross-validated myeloarchitectonic features through associations with qT1 mapping (Edwards et al., 2018; Mancini et al., 2020; Weiskopf et al., 2015). Compared to conventional weighted sequences, MP2RAGE-derived qT1 images are inherently uniform, theoretically free of other imaging properties like proton density or T2*, and are acknowledged as directly relating to cortical myelin content (Marques et al., 2017; Marques and Gruetter, 2013; van der Weijden et al., 2020). For qT1 sampling, we selected the 202 individuals from the Leipzig Study for Mind-Body-Emotion Interactions (LEMON) dataset (Babayan et al., 2019). Details of LEMON acquisition protocols and preprocessing steps have been described in detail (Mendes et al., 2019).

To correlate the depth of profiles acquired from digitized histological data microphotographs with *in vivo* intracortical qT1, we positioned 10 equivolume surfaces between the inner and outer cortical interface using *equivolumetric_surfaces.py* (https://github.com/kwagstyl/surface_tools.git). These surfaces systematically sampled the axis perpendicular to the cortical ribbon, with interpolation at each vertex (Hong et al., 2017, 2016).

### Correlation of myelin content with von Economo-Koskinas cytoarchitectonic data

To verify the neurobiological significance of MYATLAS, we correlated the MGL with von Economo-Koskinas’ cytoarchitectonic features of *gyral dome thickness*, *cellular density* and *cell size* (Scholtens et al., 2018, 2015). Since *gyral dome thickness* is reported as a range, region-specific averages were calculated. *Cellular densities* were averaged across all cortical layers, whereas *cell size* was calculated according to [H_mean_ x W_mean_]; with H_mean_ (Height) = [H_(min-max)_/2] and W_mean_ (Width) = [W_(min-max)_/2] per individual cortical layer and then averaged across all layers. To allow for between-atlas correlation, we matched the boundary of parcels using a winner-takes-all approach to assign each parcel of the von Economo-Koskinas’ atlas to our parcellation.

### Use of the MYATLAS and associated features

All parcellations and MGL maps are available in two widely used formats (gifti and nifti [‘*dlabel*’ for MGL and ‘*dscalar*’ for parcellation]) and two brain spaces (Colin27 and Conte69, both with 32k vertices). We further provide the original, manual parcel translation file (‘rh.vogt_vogt.annot’) for use with FreeSurfer. Finally, to facilitate implementation, we also provide Bash scripts which convert the original labels from the MNI-Colin27 brain to single-subject space using FreeSurfer ‘mri_label2label’ (*mapping_colin27_labels_onto_individuals[_batch].sh*). Finally, we generated a flipped version of the left hemisphere atlas based on symmetric hemispheric registration (‘*xhemf*’ command in FreeSurfer).

## Results

### 3D myeloarchitectonic atlas

The MGL patterns were similar between the MYATLAS and the Nieuwenhuys map **(Figure 1)**. Indeed, the lowest MGL values were found in highly myelinated primary sensory areas (somatosensory areas 67, 69–71II of the postcentral gyrus; auditory cortex, areas 145–157) and primary motor cortices (areas 39, 42, 43). Notably, similar to Vogt’s observation of continuous changes of architectural features, our 3D map displayed gradually decreasing myelin content in areas distant from primary cortices. Reflecting this pattern, higher-order areas (including areas 49-51 of the frontal pole) and precuneus (areas 81-85) revealed higher MGL (*i.e*., less myelination), compared to the rest of the brain. There were some noteworthy exceptions to this pattern of hierarchy-dependent myelin profiles, previously recognized by Vogt, with densely myelinated clusters, comprising the orbitofrontal (60 and 61), intraparietal (86, 87), and posterolateral (169-172) and basal (173-177, 179-180) temporal areas. These clusters have also been consistently identified both *ex vivo* and on structural MRI (Glasser and Van Essen, 2011; Nieuwenhuys and Broere, 2017), all relating to visual processing (Nieuwenhuys and Broere, 2017). Notably, areas 173,174 177,179-80 correspond to two newly discovered distinct cytoarchitectonic areas FG3 and FG4 of the fusiform gyrus (Lorenz et al., 2017).

### Intracortical depth profiling based on Vogt-Vogt classifications

**Figure 2A-B** illustrate 3D maps of fiber penetration patterns derived from Vogt-Hopf studies stratified with respect to the presence of the bands of Baillarger and bundle intrusion types (see **Method** ‘*prior work*’ for stratification details). Specifically, within available areas, high-order cognitive regions showed either the *unistriate* (temporal and frontal areas) or *bistriate* (parietal) subtype. In contrast, the bundle intrusion types were relatively homogeneous, with majority of the areas containing the *euradiate* subtype. **Figure 2C** shows patterns of myelination density across lobes.

### Correlation between surface-mapped MGL and in vivo myelin metrics

We found a positive correlation between our histology-derived MGL and *in vivo* qT1 values (p<5×10^-7^, r=0.39), indicating close correspondence between these surrogate metrics of myelin. **Figure 3A** illustrates the group-averaged qT1 map of 202 subjects selected from the LEMON dataset (Babayan et al., 2019). The lowest qT1 values were found in heavily myelinated primary cortices, with a decrease when moving towards higher order areas. Notably, low qT1 values found in the posterolateral temporal lobe likely indicate the dense myelination of the dark cluster described by Vogt (Nieuwenhuys and Broere, 2017). **Figure 3B** illustrates the correspondence between photodensitometric quantifications of cortical myelin content and *in vivo* qT1 depth profiles.

**Figure 3.**
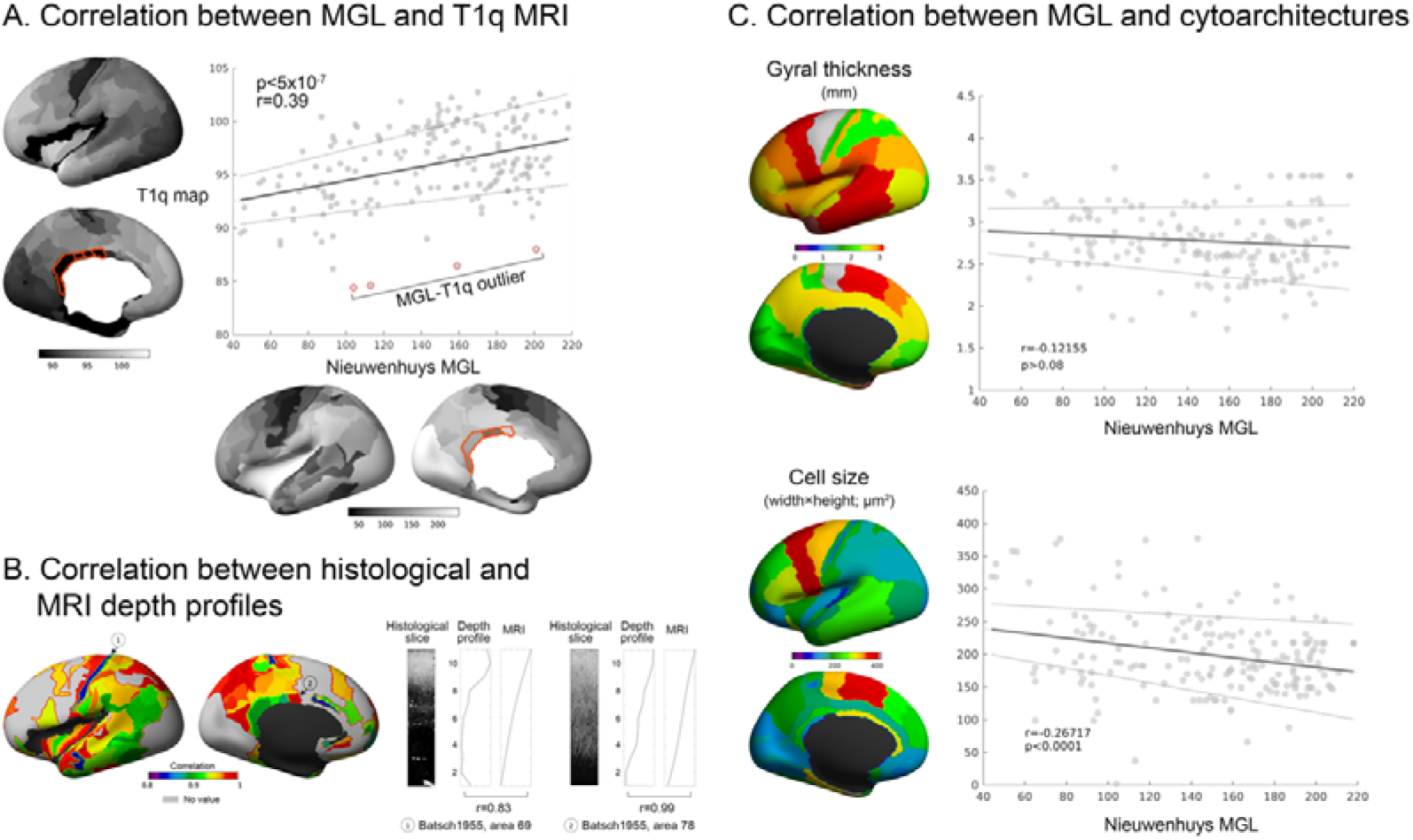
Correlation of *in vivo* MRI markers and cytoarchitectural features. **A)** Whole-brain spatial correlation between *in vivo* quantitative T1w MRI (T1q) and mean gray level (MGL) index. Outliers (>±2SD qT1) in the posterior cingulate are in red). **B)** Whole-brain depth profile correlation between qT1 and photodensitometry-derived cortical myelin content; only areas with available depth profiles are presented. Two areas with the highest and lowest correlation between histology and qT1 MRI are given as examples. **C)** Whole-brain spatial correlation between MGL and cytoarchitectural features derived from von Economo-Koskinas literature (*i.e*., gyral dome thickness and cell size).

### Correlation between surface-mapped MGL and cytoarchitectonic features

Only cell size was found to correlate with myelin density as represented by MGL (r = −0.27, p < 0.0001; **Figure 3C**), where smaller cell sizes were associated with higher myelin density. While *gyral dome thickness* trended towards a similar relationship (r = −0.12, p < 0.09), *cell density* did not (r = 0.03, p > 0.7). These findings are in line with previous studies showing that axons of smaller sensory neurons are often unmyelinated (Lee et al., 1986, p. 198). Notably, pyramidal axons of the central motor cortex exhibit complex myelination patterns (Micheva et al., 2016), which differ between cortical layers (Tomassy et al., 2014). Thus, it is conceivable that these complex interactions might not be captured by the limited resolution of a simplified metric such as MGL.

### Code and Data availability

The MYATLAS, lookup tables, source codes and the scripts for applying the atlas to Colin27 and Conte69 brain templates as well as to individual brains (“*mapping colin27_labels_onto_individuals_balch.sh*”) are available from this link (https://bic.mni.mcgill.ca/~noel/noel-myelin). A README file together with detailed descriptions of the downloadable files are available from the same web repository. All imaging-derived files are in FreeSurfer MGH and NIFTI formats and can be viewed with standard software (*e.g*., FreeView, FSLeyes or wb_view). The scripts used for data processing are available from the authors upon request.

## Discussion

Digital reconstructions of histological brain atlases constitute an important resource for contemporary neuroimaging. Such reconstructions expand availability of previously inaccessible, yet highly comprehensive, observations. Building upon the Vogt legacy, we present the MYATLAS, a 3D myeloarchitectonic digital cartography to assist neuroimaging mapping studies. To facilitate broad application, we provide the atlas together with the codes and data files. Moreover, to mitigate inter-individual variability, the parcellations and MGL maps are available both on a single- and a multi-subject group templates in stereotaxic space. Integration of detailed, quantitative data on cortical myelination will allow future neuroimaging research to assess their findings based on both myelin density and microstructure, enhancing biological validity.

Variations in myelination relate to various aspects of neocortical structure and function, including connectivity and hierarchical processing (Boshkovski et al., 2021; Huntenburg et al., 2017; Royer et al., 2020). Moreover, disrupted myeloarchitectural properties may reflect the pathological underpinning of neurological disorders (Nord et al., 2019). Thus, classification of cortical myeloarchitectonic areas and patterns through parcellation and MGL mapping has significant translational potential in both health and disease. Future neuroimaging studies may leverage this information to elucidate pathological whole-brain myeloarchitectural patterns, for instance in multiple sclerosis (Rahmanzadeh et al., 2021) and epilepsy (de Curtis et al., 2021; Drenthen et al., 2019).

Our atlas aggregates histology-derived myeloarchitectural information, depth-dependent photometric density, as well as corresponding *in vivo* qT1 profiles, together with myeloarchitectural subtypes categorized into laminar and depth intrusion patterns of fiber bundles. Notably, the high congruence between histology-derived quantifications of myelin density trough MGL and qT1 lends further biological validity to this *in vivo* microarchitectural surrogate metric easily implementable in clinical settings (Hogan, 2017; Waehnert et al., 2016). Notably, recent developments in advanced imaging sequences, such as myelin water imaging (van der Weijden et al., 2020) or high-field laminar fMRI (Trampel et al., 2019), allow for an increasingly detailed study of cortical myelin contents. In this regard, our histology-validated depth profiles can be harnessed as ground-truth data to validate future *in vivo* imaging studies investigating myeloarchitecture (O’Muircheartaigh et al., 2019; Yuan et al., 2021).

A few noteworthy points should be considered when applying MYATLAS to new data. First, as this atlas is based on consensus evidence (*i.e*., cortical boundaries) from several studies and does not incorporate information on inter-individual variability of myeloarchitectural characteristics. This limitation may however be mitigated by employing our atlas in conjunction with probabilistic cortical mapping approaches, such as *Julich Brain* (Amunts et al., 2020). Additionally, our digital atlas does not contain information on potential left-right asymmetries, since the original sources also do not contain any lateralization information. Nevertheless, a recent quantitative MRI study investigating myeloarchitectural metrics of the language system revealed heterogenous lateralization patterns (Yuan et al., 2021): While inferior frontal areas were left lateralized, the middle and superior temporal gyrus (Heschl’s gyrus and planum temporale) was found to be right lateralized. As such, future research should therefore be directed at potential functional implications of myeloarchitectural lateralization patterns in larger cohorts. Additionally, since measures of myelination density (namely MGL and cortical depth profiles) are unavailable for the occipital lobe and the insula, a whole-brain neuroimaging correlation remains somewhat partial. However, due to their high congruence, this limitation could be resolved by extrapolating MGL from qT1 *in vivo* data, preferably acquired at ultra-high magnetic field strengths (Sengupta et al., 2018).

Future applications of our architectonic mapping framework may include correlations between myeloarchitectonic density and myelin-related genes (Donkels et al., 2020; Glasser et al., 2016). For instance, building on our procedures from MGL-qT1 cross-correlation, it is now possible to relate myelin-related gene expression to myeloarchitectonic features within individual cortical parcellations. Such efforts may provide further insights on the role of specific genes in health and disease (Rahmanzadeh et al., 2021; Sprooten et al., 2019).

## CRediT author roles

- Niels A. Foit: data curation, methodology, formal analysis, interpretation of data, manuscript writing and editing.
- Seles Yung: structure segmentation, formal analysis
- Hyo-Min Lee: manuscript reviewing and editing
- Andrea Bernasconi: supervision, methodology, interpretation of data, manuscript reviewing and editing.
- Neda Bernasconi: supervision, methodology, interpretation of data, manuscript reviewing and editing
- Seok-Jun Hong: data curation, supervision, methodology, formal analysis, manuscript writing, reviewing and editing

## Declaration of interests

The authors declare that there is no conflict of interest related to this manuscript.

---

**Supplementary Table 1.**
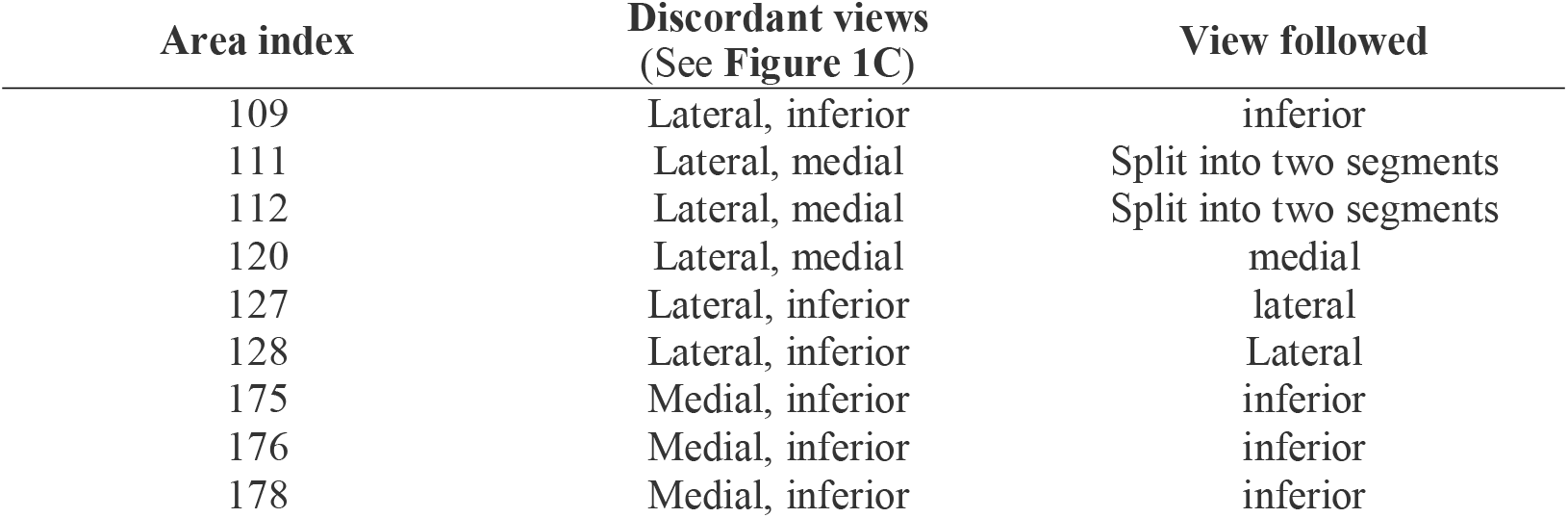
Label indices for discordant views across brain areas.

| Area index | Discordant views<br>(See Figure 1C) | View followed |
| --- | --- | --- |
| 109 | Lateral, inferior | inferior |
| 111 | Lateral, medial | Split into two segments |
| 112 | Lateral, medial | Split into two segments |
| 120 | Lateral, medial | medial |
| 127 | Lateral, inferior | lateral |
| 128 | Lateral, inferior | Lateral |
| 175 | Medial, inferior | inferior |
| 176 | Medial, inferior | inferior |
| 178 | Medial, inferior | inferior |

**Supplementary Table 2.**
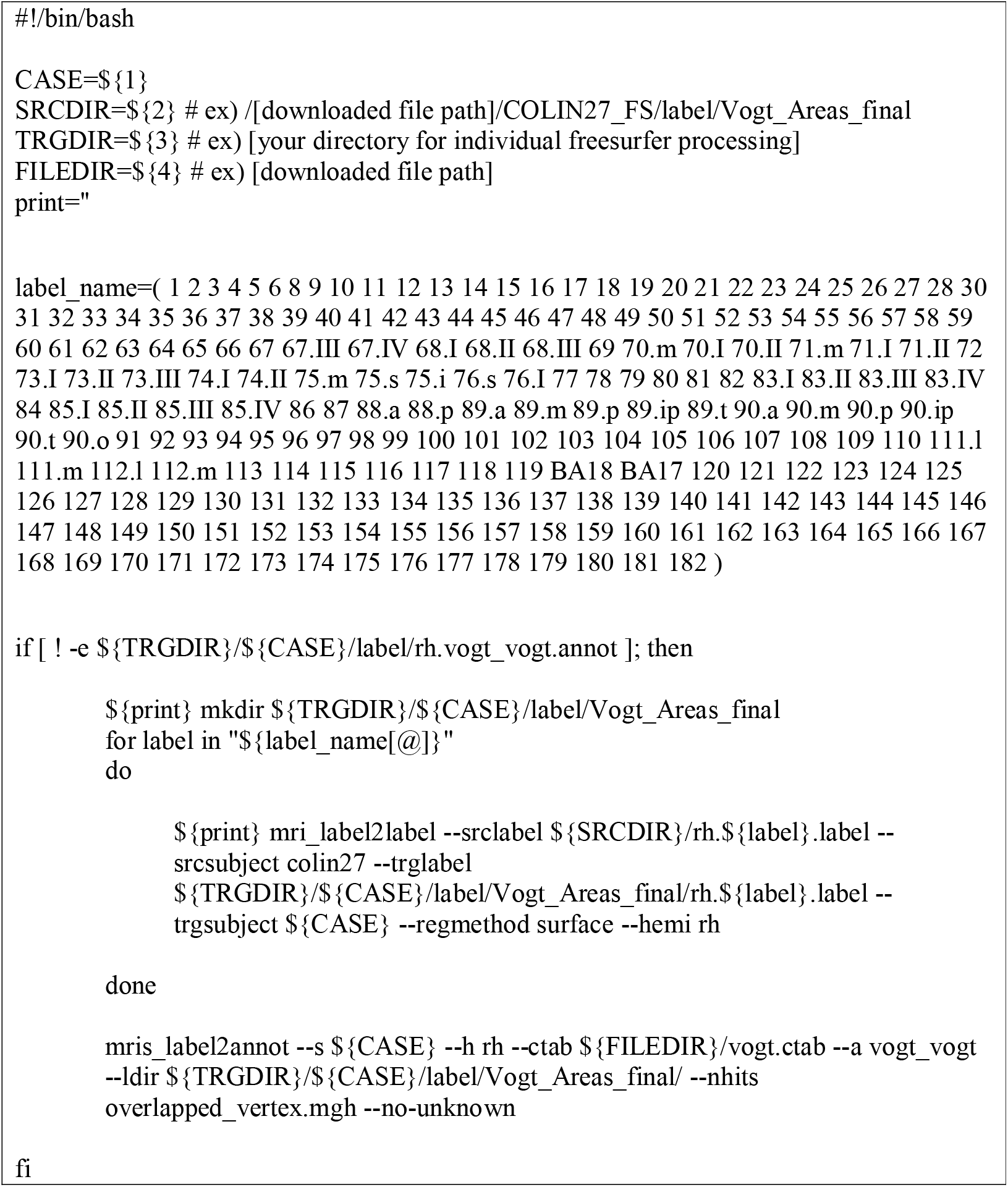
FreeSurfer command line code.

## Notes

### Competing Interest Statement

The authors have declared no competing interest.

